# Fasting and re-feeding independently alter mouse gut microbiota during intermittent fasting

**DOI:** 10.64898/2026.02.25.707984

**Authors:** Laura D. Schell, Yi Jia Liow, Rachel N. Carmody

**Author notes:** Correspondence: Laura D. Schell, Rachel N. Carmody.

## Abstract

Intermittent fasting (IF) elicits metabolic benefits that are partially driven by the gut microbiome. Studies have focused on endpoint IF-induced changes in the gut microbiome but have not explored whether the oscillating nature of IF elicits day-to-day microbiome changes that could independently affect health. To discriminate the long-term and short-term effects of IF on the gut microbiota, we fasted mice every other day (IF1:1) or every two days (IF1:2), measuring daily changes in body mass and composition, food intake, and gut microbiota composition. We show that short-term effects of fasting and re-feeding on gut microbiota composition outweigh longer-term effects of IF treatment, with composition responding differently to re-feeding and fasting. Re-feeding specifically promoted rapid expansion of *Lactobacillus*, a bacterial genus linked mechanistically to the metabolic benefits of IF. Our results highlight the plasticity of the gut microbiota, especially re-feeding effects, as a potential contributor to microbiome-mediated metabolic benefits of IF.

## INTRODUCTION

Intermittent fasting (IF) is known for its metabolic benefits in both humans and animal models, including weight loss, improved glucose homeostasis, and reduced inflammation^1^. The metabolic effects of IF have been most commonly linked to reductions in calorie intake, regulation of circadian rhythms, and the physiological impact of extended fasting periods (i.e., activation of repair mechanisms and reduced oxidative stress)^2,3^. However, recent studies suggest that IF-induced changes in the gut microbiome can also contribute causally to the metabolic benefits^4^. In a mouse model of every-other-day fasting, mice expended more energy on fed days, experienced beiging of white adipose tissue, higher body temperature, and ultimately reduced body fat^4^. Importantly, IF altered the gut microbiota, and transplantation of the IF-exposed microbiota into obese mice was capable of inducing similar metabolic improvements even without direct IF treatment. Understanding gut microbial responses to IF may thus enable us to more effectively harness the therapeutic benefits of IF.

The gut microbiota responds rapidly to dietary changes^5,6^ and has been shown to regulate diverse aspects of host energy metabolism, including appetite, nutrient absorption, energy allocation, and inflammation^7,8^. Numerous studies have found that IF alters the gut microbiota in animals subjected to every-other-day fasting, as well as in humans under popular IF interventions such as time-restricted eating (daily 12-18h fast), religious fasting, and 5:2 fasting (2 days/week of minimal calorie intake)^9,10^. Outcomes vary across studies and species, but common changes under IF include increases in microbial community richness and the relative abundance *Lactobacillus*^11–14^.

However, most studies to date have focused on endpoint changes to the gut microbiota after weeks to months of IF treatment, overlooking the fact that IF is inherently dynamic, involving frequent oscillation between the fasted and fed states. This volatility distinguishes IF from other dietary interventions known to alter the gut microbiota through the static addition or subtraction of nutrients (e.g., high-fat diets^6^, caloric restriction^15^). In the short-term, both stages of an IF cycle—fasting and subsequent compensatory overeating during refeeding—could be expected to influence the gut microbiota by affecting nutrient flows within the gut lumen^16^. In the longer-term, the intrinsic volatility of the gut environment under IF and the direct impacts of IF on host metabolism would be expected to exert their own selection pressures on the gut microbiota.

Here, we sought to distinguish the short- and long-term impacts of IF on the gut microbiota. We exposed 6-week-old chow-fed male C57BL/6J mice to every-other-day fasting (IF1:1), every-two-day fasting (IF1:2) or *ad libitum* feeding (AL) for 30 days (**Figure 1A**). Body mass and food intake were measured daily; body composition was measured at baseline and endpoint; and fecal samples were collected for gut microbial profiling on 3 baseline days and then the first 10 and last 10 days of treatment. Our data show that fasting and re-feeding exert dynamic effects on the gut microbiota that differ from each other and from longer-term microbial signatures of IF. In particular, we found that re-feeding had a greater impact on the gut microbiota than fasting, with re-feeding promoting rapid expansion of *Lactobacillus*. The varying short- and long-term effects of IF on the gut microbiota may influence host metabolism in different ways, suggesting these dynamics could potentially be exploited to optimize the metabolic response to IF.

**Figure 1.**
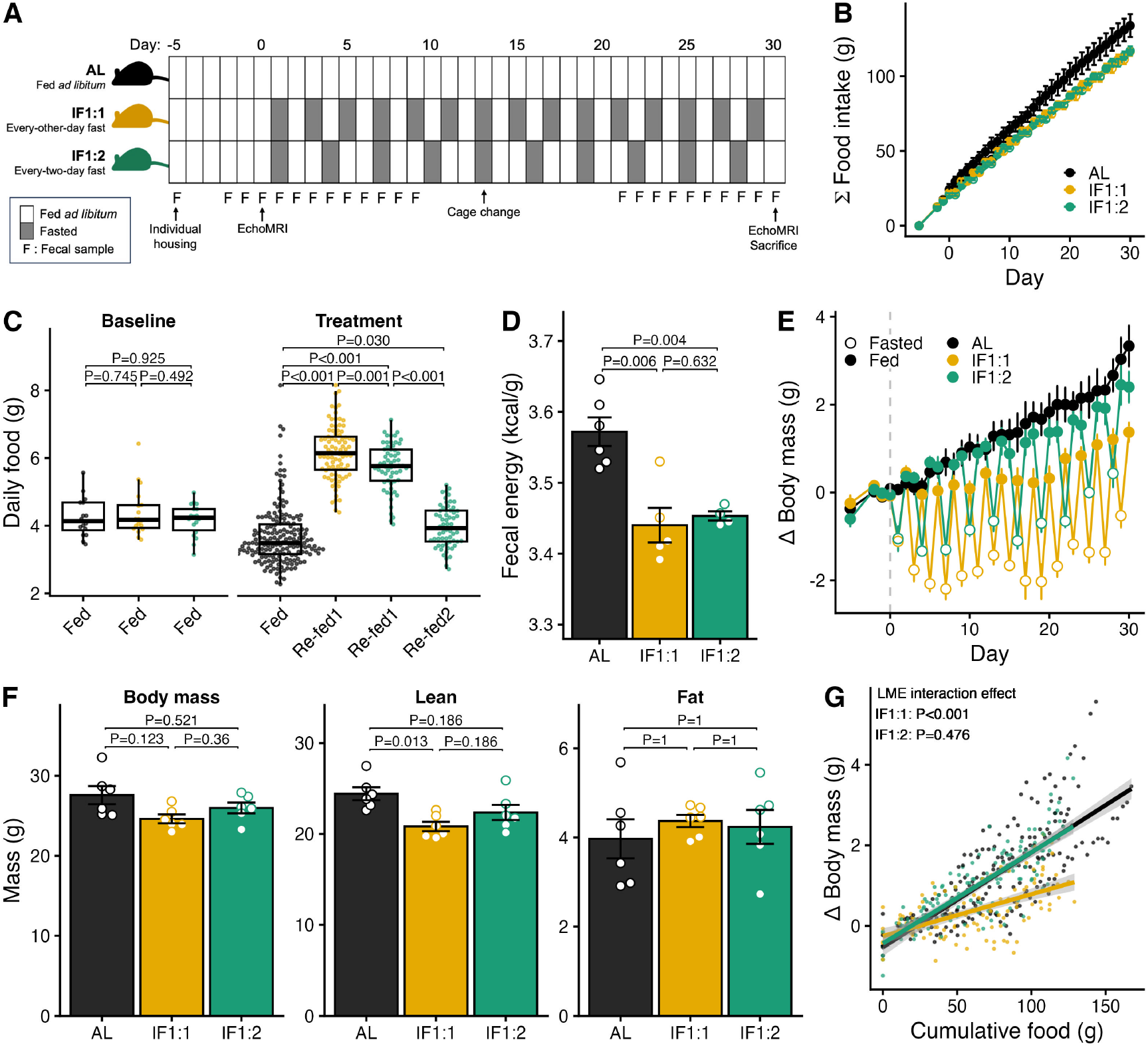
Energy balance under intermittent fasting. (A) Experimental design, n=6 mice/group. (B) Cumulative food intake from baseline through day 30 of treatment. Mean ± SEM. (C) Daily food intake for each mouse from baseline through day 30 of treatment. Stats are linear mixed effects (LME, fixed effects: feeding status; random effects: mouse ID, day). Daily food intake for AL mice was lower during treatment versus baseline (LME, fixed effects = phase; random effects = mouse ID, P_phase_=0.005), a result attributable to slowed growth rates as mice aged. Median ± IQR. (D) Energy density of feces produced from days 16-30, measured via bomb calorimetry. Stats are Student’s t-test. Mean ± SEM. (E) Change in body mass relative to day 0. Mean ± SEM. (F) Endpoint (day 30) body mass, lean mass, and fat mass, measured via EchoMRI. Stats are Student’s t-test. Mean ± SEM. (G) Change in body mass as a function of cumulative food intake, excluding fasted days. Stats are LME (fixed effects: cumulative food intake × treatment; random effects: mouse ID, day).

## RESULTS

### Intermittent fasting reduces weight gain and total food intake despite compensatory overeating and lower calorie excretion

Both groups of IF mice consumed marginally less total food than AL mice over the course of the experiment (Wilcoxon rank-sum test, P=0.052) (**Figure 1B**). The impact of fasting on total food intake was partially offset by compensatory overeating by IF mice on the first day of re-feeding after fasting (re-fed1), when IF1:1 and IF1:2 mice ate an average of 68% and 54% more food than AL mice, respectively, according to linear mixed effects models (LME) using both mouse ID and experimental day as random effects (**Figure 1C**). On the second day after fasting (re-fed2), IF1:2 mice still consumed more food than AL controls, but by a more modest 9% on average.

Over the last two weeks of treatment, IF mice also excreted 3.7% (IF1:1) and 3.3% (IF1:2) fewer calories in feces than AL controls (**Figure 1D**) suggesting greater fractional energy absorption. Overall, IF mice compensated for the reduced energy intake of fasting days by overeating on re-feeding days and excreting fewer calories via feces.

All mice gained weight over the course of treatment, but IF mice exhibited sharp decreases in weight on fasted days (**Figure 1E**). By day 30, IF1:1, but not IF1:2, were significantly lighter than AL mice, and these differences in body mass were attributable to reduced accumulation of lean mass rather than fat mass (**Figure 1F**). The absence of fat loss in IF mice was likely a result of their low initial fat mass (**Figure S1A-B**), since we conducted these experiments in healthy mice unlike many other studies focused on effects of IF in mice with obesity due to high-fat diet exposure or genetic predisposition^9^.

We next evaluated the extent to which changes in body mass could be explained by reduced food intake among IF mice using LMEs. Whereas IF1:2 and AL mice on fed days gained similar amounts of body mass for a given food intake, IF1:1 mice gained 56% less body mass than expected for their food intake (**Figure 1G**). Thus the more intensive IF1:1 treatment had an exaggerated impact on host energy balance.

### Short-term feeding oscillations are the strongest determinant of changes to gut microbiota under IF

IF-induced alterations to gut microbial composition were detectable as early as the second cycle of fasting and re-feeding (PERMANOVA, P=0.002, R^2^=0.16) and persisted through the end of the study (**Figure 2A, Table S1**), despite strong baseline patterning of microbiota composition by original source cage (PERMANOVA, P=0.001, R^2^=0.56) (**Figure S1C**). IF-induced effects on the gut microbiota were evident during both fasting and re-fed1 days (**Table S1**), and fasted days were compositionally distinct from re-fed days (PERMANOVA, P=0.001, R^2^=0.25).

**Figure 2.**
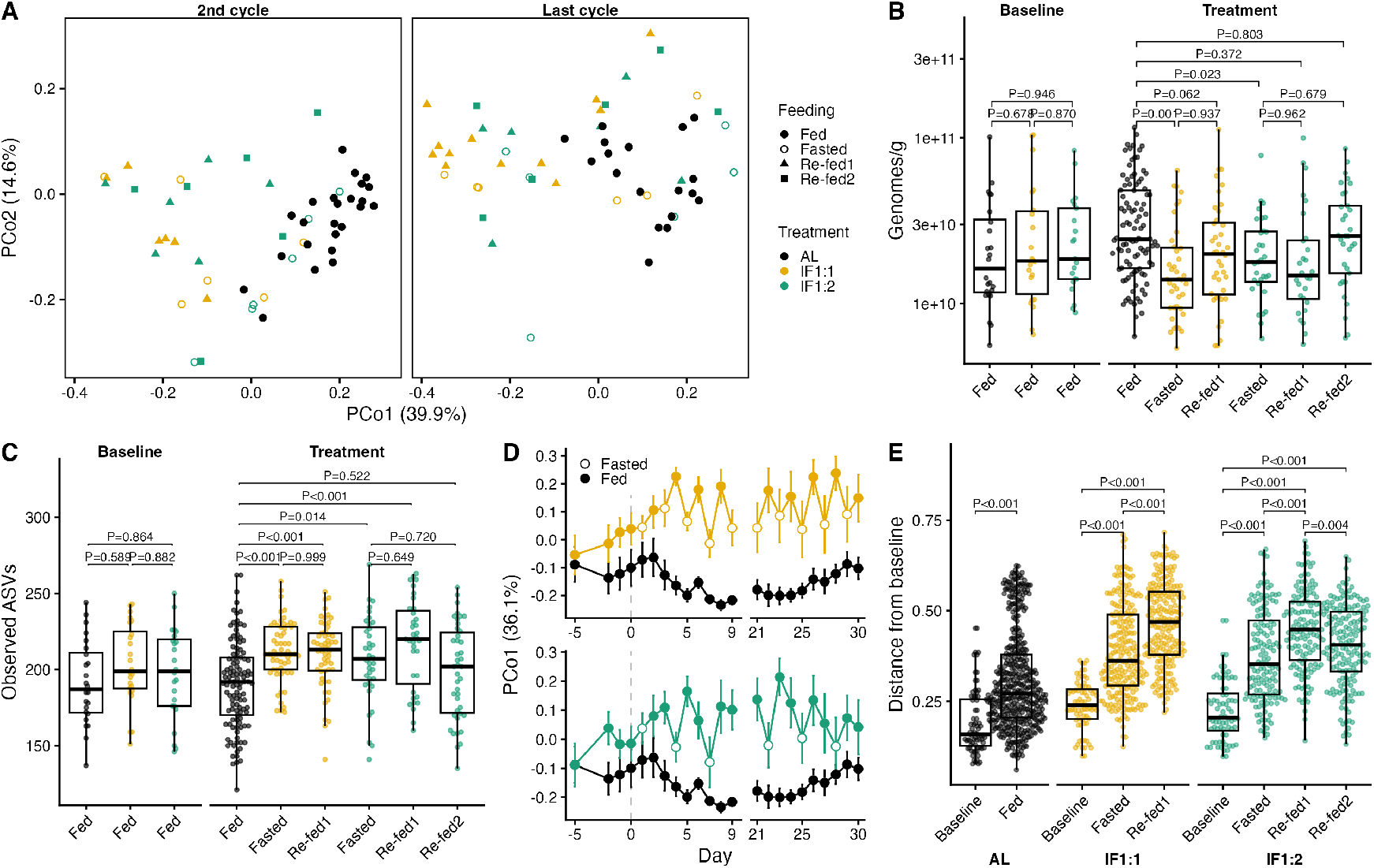
Short-term impacts of fasting and re-feeding on the gut microbiota. (A) Principal coordinate plot of Bray-Curtis distance between fecal microbiota samples during the second and final IF cycles. (B) Absolute bacterial density estimated via universal bacterial 16S qPCR. Stats are from LME model (fixed effect: feeding status; random effects: mouse ID, day). Median ± IQR. (C) Richness of ASVs per sample. Stats are from LME model (fixed effect: feeding status; random effects: mouse ID, day). Median ± IQR. (D) Bray-Curtis principal coordinate 1 (PCo1) as a function of time, showing IF1:1 versus AL (top) and IF1:2 versus AL (bottom). Mean ± SEM. (E) Bray-Curtis distance between each mouse and its own baseline days for all baseline, fasted, and fed days. Stats are from LME model (fixed effects: treatment x feeding status; random effects: mouse ID, baseline sample ID). Median ± IQR.

To distinguish between the short-term impact of feeding status and long-term impacts of IF treatment, we applied PERMANOVA/adonis tests to the fecal microbiota in the last week of treatments. The order-dependent nature of the PERMANOVA/adonis test partitions variance based on the order of variable entry, enabling us to examine the incremental effects of short-term feeding status given long-term treatment, and vice versa. These analyses showed that 27% of overall variation in the gut microbiota was attributable to IF, of which 2% was ascribable solely to the long-term effects of treatment, 8% solely to the short-term effects of feeding status, and 17% to shared (undistinguishable) effects (feeding + treatment: R^2^_feeding_ = 0.25, R^2^_treatment_ = 0.02; treatment + feeding: R^2^_feeding_ = 0.08, R^2^_treatment_ = 0.19) (**Table S1**). Thus feeding status was a stronger determinant of microbiota composition than IF treatment itself.

Both IF treatments reduced absolute bacterial abundance in the gut relative to AL feeding, particularly on fasted days (**Figure 2B**). This reduction in absolute bacterial abundance is consistent with predictions based on ecological first principles^17^ and reports of similar reductions in humans on very-low-calorie diets^15^. Despite IF-induced reductions in absolute bacterial abundance, within-sample taxonomic richness (alpha-diversity) increased among IF1:1 and IF1:2 mice on fasted and re-fed1 days compared to both baseline and AL-fed controls (**Figure 2C**). Enhanced competition between microbes due to the volatility of nutrient availability under IF may promote taxonomic richness, or else reductions in the absolute abundance of most taxa may allow for less sensitive, low-abundance taxa to reach detectable levels.

IF treatment therefore altered the composition, abundance, and diversity of the gut microbiota, although the type and magnitude of these changes differed by feeding status.

### Gut bacteria respond to both short-term cycling and long-term treatment patterns

Different bacterial taxa were independently sensitive to the short-term and long-term features of IF treatment. Analysis of differentially abundant taxa using MaAsLin3^18^ identified 55 genera that were significantly enriched or depleted during fasting and/or re-feeding relative to baseline and AL mice (**Figure 3, Table S2**).

**Figure 3.**
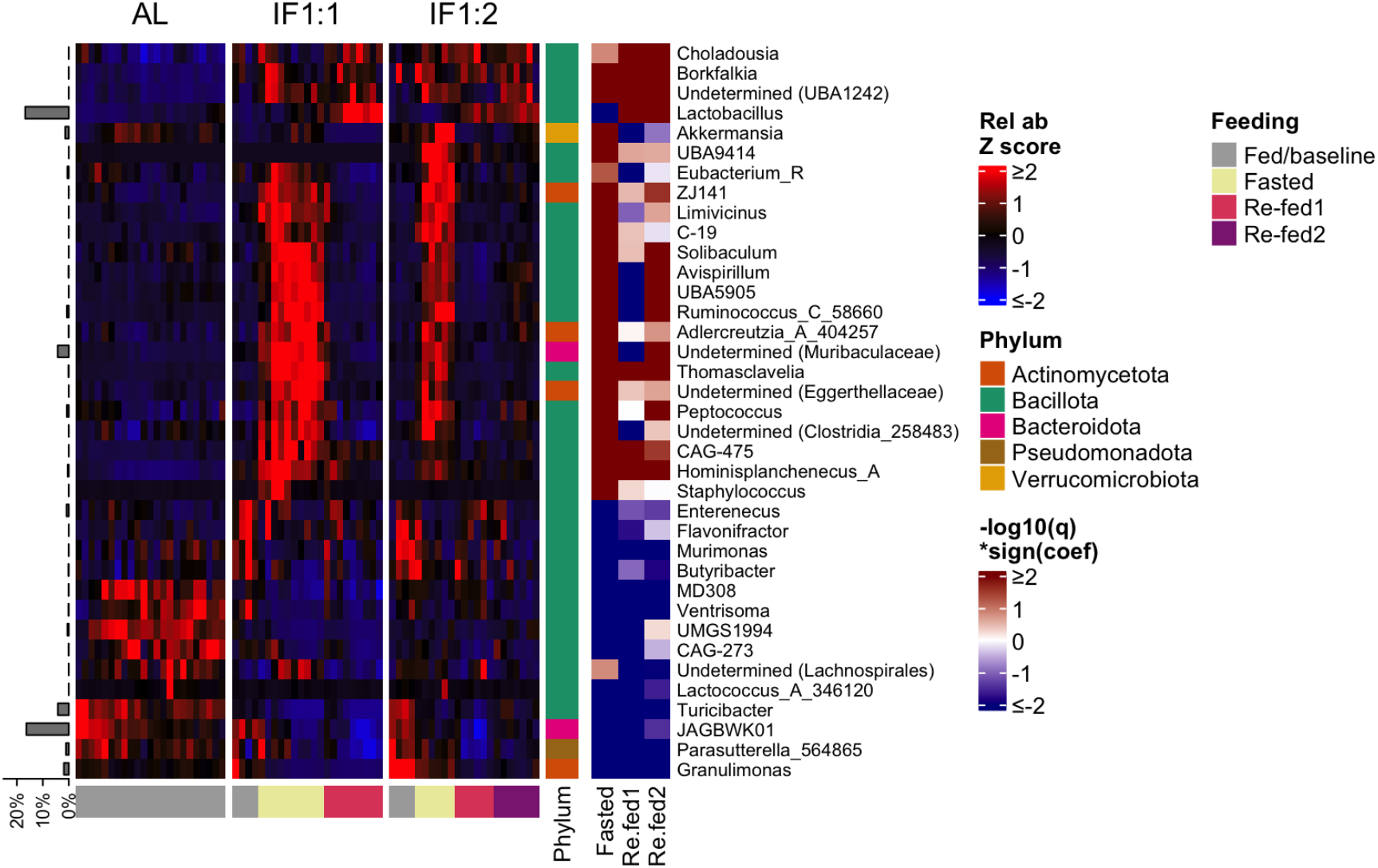
Bacterial genera significantly associated with fasting and/or re-feeding. Rows are genera that were differentially abundant in feces across feeding status, identified using a MaAsLin3 linear mixed effects model (fixed effects: feeding; random effects: mouse ID, source cage ID, day) and remained significant under both the standard unscaled model and when scaling relative abundances by per-sample absolute bacterial abundance. Where genus name is undetermined, the next lowest known taxonomic classification is indicated. Columns represent individual days, ordered by treatment group and feeding status then chronologically. Data are row z-scores of mean relative abundance by genus. Annotations: mean genus relative abundance across all samples (left); feeding status (bottom); phylum categorization of each genus (inner right); significance and direction of feeding effects on genus relative abundance identified by the unscaled MaAsLin3 model (right).

Some genera responded primarily to either fasting or re-feeding, without longer-term changes attributable to IF treatment. Fasting days were characterized by increases in the relative abundance of many genera, including *Ruminococcus, Akkermansia, Thomasclavelia* and 12 others that were elevated during fasting but not re-feeding (**Figure 3, 4A-C, S3A**). By contrast, re-feeding was characterized largely by an expansive bloom in *Lactobacillus*, which reached a mean relative abundance of >30% in both IF1:1 and IF1:2 mice on some re-fed1 days (**Figure 4D**). Six genera that displayed elevated relative abundance during fasting were also significantly reduced during re-feeding (**Figure 3**).

**Figure 4.**
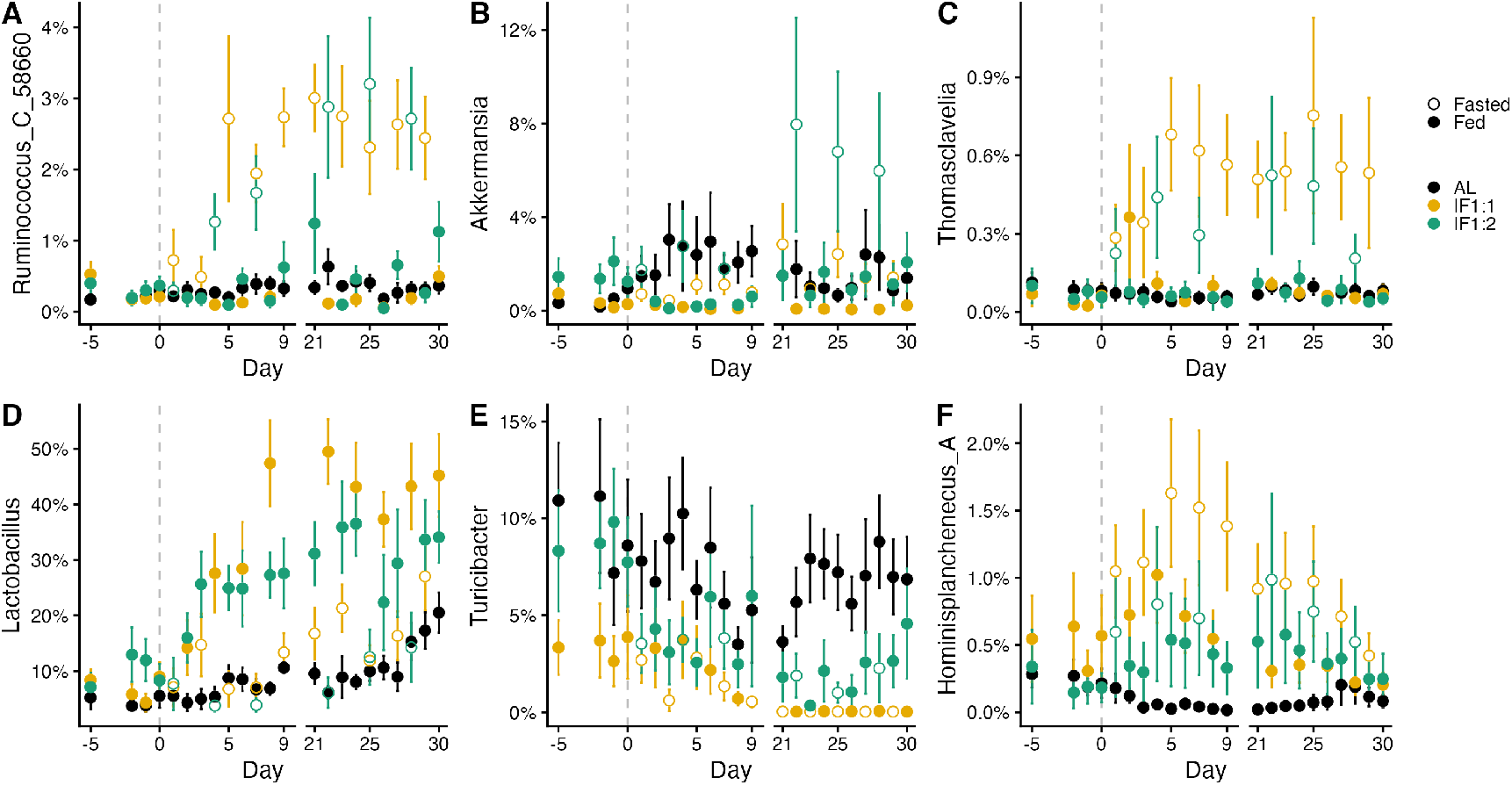
Examples of differential responses of gut microbial genera to the long- and short-term impacts of intermittent fasting. (A) Relative abundance of *Ruminococcus* is elevated only during fasting (fasted: β=2.51, q=1.08×10^-8^). (B) Relative abundance of *Akkermansia* is elevated during fasting and reduced during re-feeding (fasted: β=1.82, q=7.59×10^-4^; re-fed1: β= -1.55, q=1.94×10^-5^). (C) Relative abundance of *Thomasclavelia* is elevated only during fasting (fasted: β=3.68, q=4.07×10^-18^). (D) Relative abundance of *Lactobacillus* is elevated during re-feeding (re-fed1: β=1.82, q=1.23×10^-6^; re-fed2: β=1.66, q=1.23×10^-6^). (E) Relative abundance of *Turicibacter* is reduced long-term in IF mice regardless of feeding status (fasted: β= -2.77, q=2.57×10^-7^; re-fed1: β= -2.88, q=3.49×10^-7^; re-fed2: β= -2.14, q=0.006). (F) Relative abundance of *Hominisplanchenecus* is elevated in IF mice with an exacerbated increase during fasting (fasted: β=2.60, q=2.50×10^-6^; re-fed1: β=1.50, q=0.020; re-fed2: β=1.96, q=0.001). (A-F) Mean ± SEM. Statistics are from MaAsLin3 model (fixed effect: feeding status; random effects: mouse ID, source cage ID, day), not scaled to absolute bacterial abundance. β-coefficients indicate log2 fold change from baseline/AL-fed samples and the q-value is the FDR corrected P-value.

Other genera responded primarily to long-term IF treatment rather than short-term food cycling, such as *Turicibacter*, JAGBWK01 (a member of Muribacillaceae), and *Parasuterella*, all of which were depleted by IF treatment on both fasted and re-fed days (**Figure 4E, S3**). Still other genera showed both consistent long-term treatment effects and responses to daily food cycling. The genus *Hominisplanchenecus*, for instance, was elevated during all stages of the IF cycle, with a greater increase during fasting (**Figure 4F**).

Given that IF mice exhibited reduced absolute bacterial abundance on fasted days (**Figure 2B**), we sought to distinguish whether any of the significant increases in genus relative abundance during fasting represented increases in absolute abundance or were artifacts driven by absolute depletions in other taxa (**Figure S2A-B**). Bacteria that reside in the mucus lining like *Akkermansia*, for instance, might be expected to be more resistant to depletion due to absence of dietary substrates during fasting, thus showing an increase in relative abundance even if absolute abundance remained unchanged. We therefore ran MaAsLin3 after scaling bacterial relative abundance values by the absolute bacterial abundance of each sample (**Table S2**).

With absolute abundance scaling, 37 of the 55 previously identified differentially abundant genera remained differentially abundant (α = 0.05, FDR-corrected q values) (**Figure S2C**), indicating that most changes were not artifacts of reduced bacterial density. However, coefficients in the scaled models were shifted negatively relative to the unscaled coefficients (**Figure S2D**). Therefore, the scope and magnitude of these IF-associated increases in relative abundance were somewhat more modest on an absolute scale, but IF-associated reductions in relative abundance were also slightly greater in magnitude at an absolute scale.

### Attenuated gut microbiome changes during the second day of re-feeding

Given the less frequent fasting of IF1:2 mice, their lesser degree of overeating on re-fed1 and re-fed2 days, and the reduced long-term impact of IF1:2 treatment on body mass and body composition, we predicted that IF1:2 mice would experience weaker changes in the gut microbiota compared with IF1:1 mice. Differences in the gut microbiota between the two treatments were most evident on re-fed2 days, which were compositionally distinct from both re-fed1 days (PERMANOVA, P=0.001, R^2^=0.06) and fasting days (P=0.001, R^2^=0.08). Re-fed2-associated changes to the fecal microbiota were attenuated relative to re-fed1 day in terms of overall compositional distance from baseline and AL controls (**Figure 2E, S1E**) and differentially abundant microbes (**Figure 3**). Absolute bacterial abundances on re-fed2 days were indistinguishable from those of AL controls (**Figure 2B**), suggesting rapid recovery of microbial biomass after a single day of re-feeding.

Outside of re-fed2 days, gut microbiota differences between IF1:2 and IF1:1 treatment were detectable but limited in scope. Over the last week of treatment, IF1:2 microbiota differed from IF1:1 microbiota on matched fasted and refed-1 days, but these differences had relatively small effect sizes (PERMANOVA, P=0.001, R^2^=0.03) (**Table S1**). Accounting for differences by treatment group in MaAsLin3 models, we detected an attenuated increase in the relative abundance of *Thomasclavelia* among IF1:2 mice during fasting (P=0.016), but this difference did not reach significance when controlling for total bacterial abundance (P=0.429) (**Figure 4C**). Therefore, to the extent that reduced weight loss under IF1:2 versus IF1:1 treatment was linked to changes in the gut microbiota, this was likely due to reduced frequency of fasting and re-fed1 days rather than any unique impact of IF1:2 treatment on the gut microbiota.

### Re-feeding promotes Lactobacillus and drives greater changes in gut microbiota composition than fasting

Each cycle of fasting and re-feeding drove consistent oscillations in gut microbial composition, but it was the gut microbiota during re-feeding, rather than fasting, that differed most from the AL fed state. Principal coordinate analysis of Bray-Curtis distance between samples shows that gut microbiota signatures on re-fed days shifted away from AL and baseline signatures along PCo1 and PCo2, whereas fasted days were intermediate between re-fed and AL signatures (**Figure 2D, S1D**). Additionally, re-fed gut microbiota were more dissimilar than fasted gut microbiota relative to baseline (**Figure 2E**) and to day-matched AL mice (**Figure S1E**).

The main compositional shift underlying gut microbiota changes during re-feeding was a bloom in *Lactobacillus*, which reached >30% relative abundance on many re-fed1 days in both IF1:1 and IF1:2 mice and >50% relative abundance in some IF1:1 mice (**Figure 4D**). Changes in *Lactobacillus* have been previously observed in mice exposed to IF^11–14^, and the genus is notable for its rapid growth and ability to ferment carbohydrates to produce acetate and lactate^19^. During IF, the larger influx of food on re-fed1 days likely caused more nutrients to escape host digestion, potentially allowing endogenous *Lactobacillus* to expand rapidly and temporarily outcompete slower-growing taxa. Importantly, previous mouse models of IF1:1 linked metabolic improvements to gut microbial production of acetate and lactate that subsequently promoted adipocyte beiging and thermogenesis, with higher total energy expenditure on re-feeding days^4^. Acetate, lactate, *Lactobacillus*, and other commensal lactic acid bacteria have also been shown to promote brown fat thermogenesis in other contexts^20–22^. Together, these results suggest that expansion of *Lactobacillus* in response to re-feeding—but not during fasting—may underpin microbiome-mediated metabolic improvements with IF treatment. This re-feeding-focused view contrasts with conventional focus among IF researchers on the physiological impacts of extended fasting, suggesting that both may be important in eliciting the metabolic benefits of IF.

## DISCUSSION

Intermittent fasting poses a unique ecological challenge for gut microbes, given its alterations to the gut environment on both short- and long-term timescales. In our experiments involving every-other-day (IF1:1) and every-two-day (IF1:2) fasting, mice lost weight and absolute gut bacterial abundance was roughly halved on average during fasting. That fasting should impact the microbiome is unsurprising, as very-low-calorie diets and ketogenic diets that induce fasting-like physiological responses in the host have been shown to affect gut microbiome composition and function^15,23^. However, our work emphasizes that compensatory overeating during the re-feeding period may be just as important as fasting in shaping the impact of intermittent fasting on the gut microbiome.

Our study underscored that the impacts of fasting and re-feeding on the gut microbiota are distinct. On the first day of re-feeding, when IF mice consumed >50% more food than expected, the gut microbiota exhibited the starkest differences from baseline and *ad libitum*-fed controls. In both IF1:1 and IF1:2 groups, the gut microbiota on the first day of re-feeding showed a particularly large expansion of *Lactobacillus*, a bacterial genus previously found to be enriched with IF^11–14^ and causally linked to the metabolic benefits of IF via acetate and lactate production^4^. Crucially, our data suggest that elevated post-fast re-feeding, rather than fasting or IF treatment *per se*, may account for these microbiota-mediated benefits, although additional mechanistic testing is required to confirm this hypothesis.

While the IF1:1 and IF1:2 fasting models employed here are ideal for understanding the ecological impact of fasting and re-feeding cycles on the gut microbiota, such day-to-day variation will be harder to detect in humans given our substantially longer gastrointestinal transit times (12-72 hr in humans^24,25^, 2-6 hr in mice^26,27^). This would be especially true given the shorter fasting periods of time-restricted eating regimens, in which food is consumed during a 6-10-hr window each day. However, lack of sampling resolution does not preclude the possibility that such short-term changes to the gut microbiome during IF do occur in humans. Previous studies have consistently reported IF-associated changes in gut microbiota composition^9,10^, although the inability to reliably link samples to either the fasted or re-fed state likely adds considerable noise to experimental outcomes. New technologies for *in vivo* sampling in healthy humans^28^ will be an important tool for future translational research in this area.

In conclusion, our study showed that, during IF treatment, the gut microbiota responds uniquely to each stage of the feeding cycle, with additional changes under long-term treatment. Evidence that microbiota changes previously linked to the metabolic benefits of IF were restricted to the re-feeding stage complements current research emphasis on fasting and suggests that both stages may offer important levers for maximizing health benefits. Further research is necessary to establish this pattern in humans and to determine whether the long-term impacts of IF on the microbiome result from changes to host physiology or compounding selective pressures on the gut microbiota with each additional feeding cycle.

## ACKNOWLEDGEMENTS

This work was supported by funding from NIGMS 1R35GM160067 (RNC), NSF BCS-2142073 (LDS and RNC), NSF Graduate Research Fellowship (LDS), JSPS KAKENHI Grants-in-Aid for Scientific Research 23KJ0457 (YJL), and the William F. Milton Fund (RNC). The authors thank our colleagues in the Nutritional and Microbial Ecology Lab for feedback on early drafts, and particularly Michelle Stegawski for her work on fecal bomb calorimetry. We also thank staff members of Harvard University’s Bauer Core Facility, Office of Animal Resources, and Biological Research Infrastructure for their assistance with sequencing and animal work, respectively. The content of this article is solely the responsibility of the authors and does not necessarily represent the official views of the National Institutes of Health.

## AUTHOR CONTRIBUTIONS

This study was conceived and designed by LDS and RNC. All animal work and sample collection was performed by LDS. Sample processing was performed by LDS and YJL. Data were analyzed by LDS. Manuscript was written by LDS and RNC with contributions and final approval of all authors.

## DATA AND CODE AVAILABILITY

The 16S rRNA gene sequencing data described in this study are available in the NCBI Sequence Read Archive under BioProject number PRJNA1428163. All other data and the code used to analyze data and generate figures will be made available upon reasonable request.

## METHODS

### Animal husbandry

Male C57BL/6J mice were individually housed under specific pathogen-free conditions in Harvard University’s Biological Research Infrastructure facility and maintained on a standard chow diet (Prolab IsoPro RMH 3000) and 12h:12h light/dark cycle for the duration of the study. Food was weighed and added or removed at ∼8 hours into the light cycle each day, and during these interventions data on body mass and fecal samples were collected. Wellness checks were performed daily to identify any potential adverse responses to treatment, and no mice were excluded from the study. All animal work was reviewed and approved by the Harvard University Institutional Animal Care and Use Committee under protocol #17-06-306.

### Body composition measurement

Lean body mass and fat mass of live animals were quantified using an EchoMRI-100H system (EchoMRI, Houston, TX).

### Quantification of fecal energy density

Cage bottoms from the last 2 weeks of the study period were collected and sifted to separate fecal material. Between 1.7-2.0 g of fecal material was combusted in a Parr 6050 Bomb Calorimeter (Parr Instrument Company, Moline, IL) to quantify caloric density.

### Fecal DNA isolation

Fresh fecal samples were collected directly from mice, immediately flash frozen in liquid nitrogen, and stored at -80ºC until processing. DNA was extracted using the E.Z.N.A Soil DNA kit (Omega BioTek, Norcross, GA) according to manufacturer instructions.

### Quantification of absolute bacterial abundance

Bacterial abundance was quantified in fecal DNA extracts via qPCR targeting the V4 region of the 16S rRNA gene, with assays performed in triplicate. To set up reactions, 2 μl template DNA was combined with 5 μl PowerUp SYBR Green Mix (2X) (ThermoFisher, Waltham, MA), 2 μl nuclease-free water, and 0.5 μl each of 10 nM unbarcoded primers (515F, 806R). Reactions were run on a BioRad CFX 96-well real-time PCR thermocycler (BioRad Laboratories, Waltham, MA) with the following protocol: enzyme activation for 2 min at 50ºC; initial denature for 2 min at 95ºC; then 40 cycles of 95ºC for 3 sec and 60ºC for 30 sec. Any replicates with a quantification threshold (Cq) ≥1 different from the sample mean were excluded, and samples with ≥2 replicates meeting these criteria were re-run. Bacterial 16S rRNA gene concentrations were calculated against a standard curve of genomic DNA isolated from a pure culture of *E. coli* and a conversion factor of 2.03 × 10^5^ (based on an estimated mean gut bacterial genome size of 4.50 Mbp)^29^ was used to calculate bacterial genome equivalents.

### 16S rRNA gene sequencing

DNA extracts were prepared for sequencing as described previously^30^. Briefly, the V4 region of the 16S rRNA gene was PCR-amplified with barcoded primers (515F, 806R), run on a gel to confirm amplification and ensure no contamination was present. PCR amplicons were then quantified using the Quant-iT Picogreen dsDNA Assay Kit (Invitrogen, Carlsbad, CA), pooled evenly according to DNA concentration, cleaned using AmpureXP beads (Agencourt, Brea, CA), then gel-purified and extracted using the Qiaquick Gel Extraction Kit (Qiagen, Hilden, Germany).

Prepared 16S libraries were sequenced 2×150 bp on one lane of an Illumina NovaSeq S1 platform, with a 30% PhiX spike-in to provide sequence diversity. Demultiplexed sequences were processed using Qiime2^31^ and denoised with Dada2^32^, truncating at 149 bp to optimize read quality. A total of 123,663,276 reads, representing 412 samples (median 291,487 reads/sample) and 1,355 unique amplicon sequence variants (ASVs), were retained after quality filtering. Sequence taxonomy was assigned using a qiime2 pre-trained classifier for the 515F/806R region based on the Greengenes2 2024.09 database^33^. Further processing was performed in R (v4.5.1) using qiime2R (v0.99.6) for the initial import and phyloseq (v1.52.0) for preprocessing and subsequent analysis. Data were pre-processed by pruning low-abundance ASVs (defined as <3 reads across all samples) and rarefied to 40,000 reads/sample for even sequencing depth, which resulted in a total of 965 ASVs being represented across our 412 samples.

### Statistical methods

All statistical tests were performed in R (v4.5.1). For cross-sectional comparisons, we applied either t-tests or Wilcoxon rank-sum tests for normal- and non-normally distributed data, respectively, as determined by the Shapiro-Wilkes test. Unless otherwise specified, all comparisons used the AL treatment group or AL-fed state (baseline and AL group) as a reference, and multiple hypothesis testing was corrected using the Holm-Bonferroni method. For analyses across the longitudinal dataset, linear mixed effects models were applied with the nlme package (v3.1-168) and comparisons between the coefficients for different variable levels were performed using Tukey’s Honest Significant Difference test via the lsmeans package (v2.30-2).

Overall microbiome composition differences were assessed using PERMANOVA/adonis2 tests via the vegan package (v2.7-2). Identification of differentially abundant genera was performed using MaAsLin3^18^ (v1.1.2), which applies linear mixed effects models to log-transformed data and corrects for multiple hypothesis testing with the false discovery rate (FDR) correction.

