## Supplemental materials for "Fasting and re-feeding independently alter mouse gut microbiota during intermittent fasting"


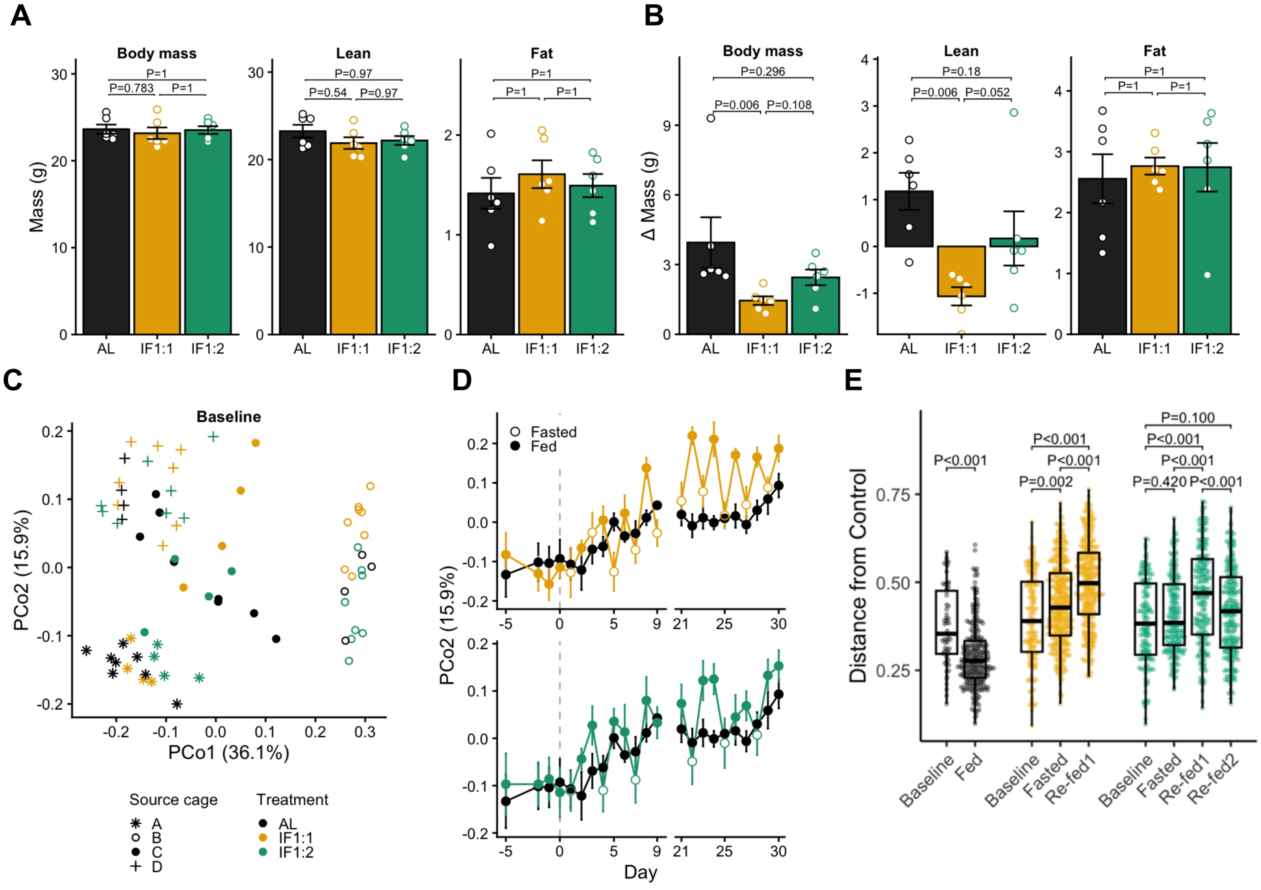


**Figure S1. Effects of intermittent fasting on energy balance and gut microbiota composition**

(A-B) Body composition at baseline (A) and change in body composition between baseline and day 30 (B). Stats are Wilcox rank-sum test. Mean ± SEM.

(C) Bray-Curtis principal coordinate plot showing baseline variation in fecal microbiome composition by source cage (days -5, -2, -1, 0).

(D) Bray-Curtis principal coordinate 2 (PCo2) as a function of time, showing IF1:1 versus AL (top) and IF1:2 versus AL (bottom). Mean ± SEM.

(E) Bray-Curtis distance between each sample and each day-matched AL sample. Stats are Wilcox rank-sum test. Median ± IQR.

### Table S1. PERMANOVA/adonis2 statistics

| Data subset | Terms | Feeding | | Treatment | | Source cage | | Day | | MouseID | |
|---|---|---|---|---|---|---|---|---|---|---|---|
| | | $R^2$ | *P* | $R^2$ | *P* | $R^2$ | *P* | $R^2$ | *P* | $R^2$ | *P* |
| Baseline: All samples | Source cage + Day | | | | | 0.559 | **0.001** | 0.011 | 0.114 | | |
| Cycle 1: AL fed vs fasted | Source cage + Feeding | 0.046 | 0.307 | | | 0.428 | **0.001** | | | | |
| Cycle 1: AL fed vs fasted | Source cage + Feeding + Treatment group | 0.046 | 0.305 | 0.065 | 0.098 | 0.428 | **0.001** | | | | |
| Cycle 1, AL fed vs re-fed1 | Source cage + Feeding | 0.064 | 0.128 | | | 0.463 | **0.001** | | | | |
| Cycle 1, AL fed vs re-fed1 | Source cage + Feeding + Treatment group | 0.064 | 0.164 | 0.006 | 0.992 | 0.463 | **0.001** | | | | |
| Cycle 1, fasted vs re-fed1 | Source cage + Feeding | 0.066 | 0.265 | | | 0.125 | 0.075 | | | | |
| Cycle 1, fasted vs re-fed1 | Source cage + MouseID + Feeding | 0.055 | 0.410 | | | 0.125 | 0.091 | | | 0.030 | 0.315 |
| Cycle 2: AL fed vs fasted | Source cage + Feeding | 0.162 | **0.002** | | | 0.166 | **0.057** | | | | |
| Cycle 2: AL fed vs fasted | Source cage + Feeding + Treatment group | 0.162 | **0.001** | 0.069 | 0.059 | 0.166 | **0.039** | | | | |
| Cycle 2: AL fed vs re-fed1 | Source cage + Feeding | 0.357 | **0.001** | | | 0.297 | **0.001** | | | | |
| Cycle 2: AL fed vs re-fed1 | Source cage + Feeding + Treatment group | 0.357 | **0.001** | 0.020 | 0.330 | 0.297 | **0.001** | | | | |
| Cycle 2: fasted vs re-fed1 | Source cage + Feeding | 0.127 | **0.001** | | | 0.230 | **0.001** | | | | |
| Cycle 2: fasted vs re-fed1 | Source cage + MouseID + Feeding | 0.132 | **0.001** | | | 0.230 | **0.001** | | | 0.040 | 0.118 |
| Final week: AL fed vs fasted | Source cage + Day + Feeding | 0.168 | **0.001** | | | 0.215 | **0.001** | 0.010 | 0.277 | | |
| Final week: AL fed vs re-fed1 | Source cage + Day + Feeding | 0.322 | **0.001** | | | 0.201 | **0.001** | 0.008 | 0.187 | | |
| Final week: fasted vs re-fed1 | Source cage + Feeding | 0.125 | **0.001** | | | 0.295 | **0.001** | | | | |
| Final week: fasted vs re-fed1 | Source cage + MouseID + Feeding | 0.125 | **0.001** | | | 0.295 | **0.001** | | | 0.015 | 0.088 |
| Final week: fasted vs re-fed1 | Source cage + Day + Feeding | 0.125 | **0.001** | | | 0.295 | **0.001** | 0.006 | 0.591 | | |
| Final week: fasted vs re-fed1 | Source cage + Day + MouseID + Feeding | 0.125 | **0.001** | | | 0.295 | **0.001** | 0.006 | 0.603 | 0.015 | 0.098 |
| Final week: All samples | Source cage + Day + Feeding + Treatment group | 0.250* | **0.001** | 0.022* | **0.002** | 0.196 | **0.001** | 0.005 | 0.225 | | |
| Final week: All samples | Source cage + Day + Treatment group + Feeding | 0.083* | **0.001** | 0.188* | **0.001** | 0.196 | **0.001** | 0.005 | 0.244 | | |
| Final week: IF1:1 vs IF1:2 (fasted) | Source cage + Treatment group | | | 0.061 | **0.035** | 0.331 | **0.001** | | | | |
| Final week: IF1:1 vs IF1:2 (fasted) | Source cage + Day + Treatment group | | | 0.059 | **0.040** | 0.331 | **0.001** | 0.006 | 0.907 | | |
| Final week: IF1:1 vs IF1:2 (re-fed1) | Source cage + Treatment group | | | 0.042 | **0.035** | 0.372 | **0.001** | | | | |
| Final week: IF1:1 vs IF1:2 (re-fed1) | Source cage + Day + Treatment group | | | 0.042 | **0.034** | 0.372 | **0.001** | 0.016 | 0.396 | | |
| Final week: IF1:1 vs IF1:2 (fasted & re-fed1) | Source cage + Feeding + Treatment group | 0.269 | **0.001** | 0.025 | **0.003** | 0.193 | **0.001** | | | | |
| Final week: IF1:1 vs IF1:2 (fasted & re-fed1) | Source cage + Day + Feeding + Treatment group | 0.268 | **0.001** | 0.025 | **0.001** | 0.193 | **0.001** | 0.005 | 0.322 | | |
| Final week: refed2 vs refed1 (IF1:2 only) | Source cage + Feeding | 0.055 | **0.002** | | | 0.374 | **0.001** | | | | |
| Final week: refed2 vs refed1 (IF1:2 only) | Source cage + MouseID + Feeding | 0.055 | **0.001** | | | 0.374 | **0.001** | | | 0.313 | **0.001** |
| Final week: refed2 vs fasted (IF1:2 only) | Source cage + Feeding | 0.082 | **0.014** | | | 0.355 | **0.001** | | | | |
| Final week: refed2 vs fasted (IF1:2 only) | Source cage + MouseID + Feeding | 0.082 | **0.001** | | | 0.355 | **0.001** | | | 0.403 | **0.001** |

* These two tests use the same terms, but the order of the two related terms Feeding and Treatment is switched. Therefore, while the two $R^2$ values for these terms sum to the same amount across both tests, any shared descriptive power is ascribed to the term listed first.

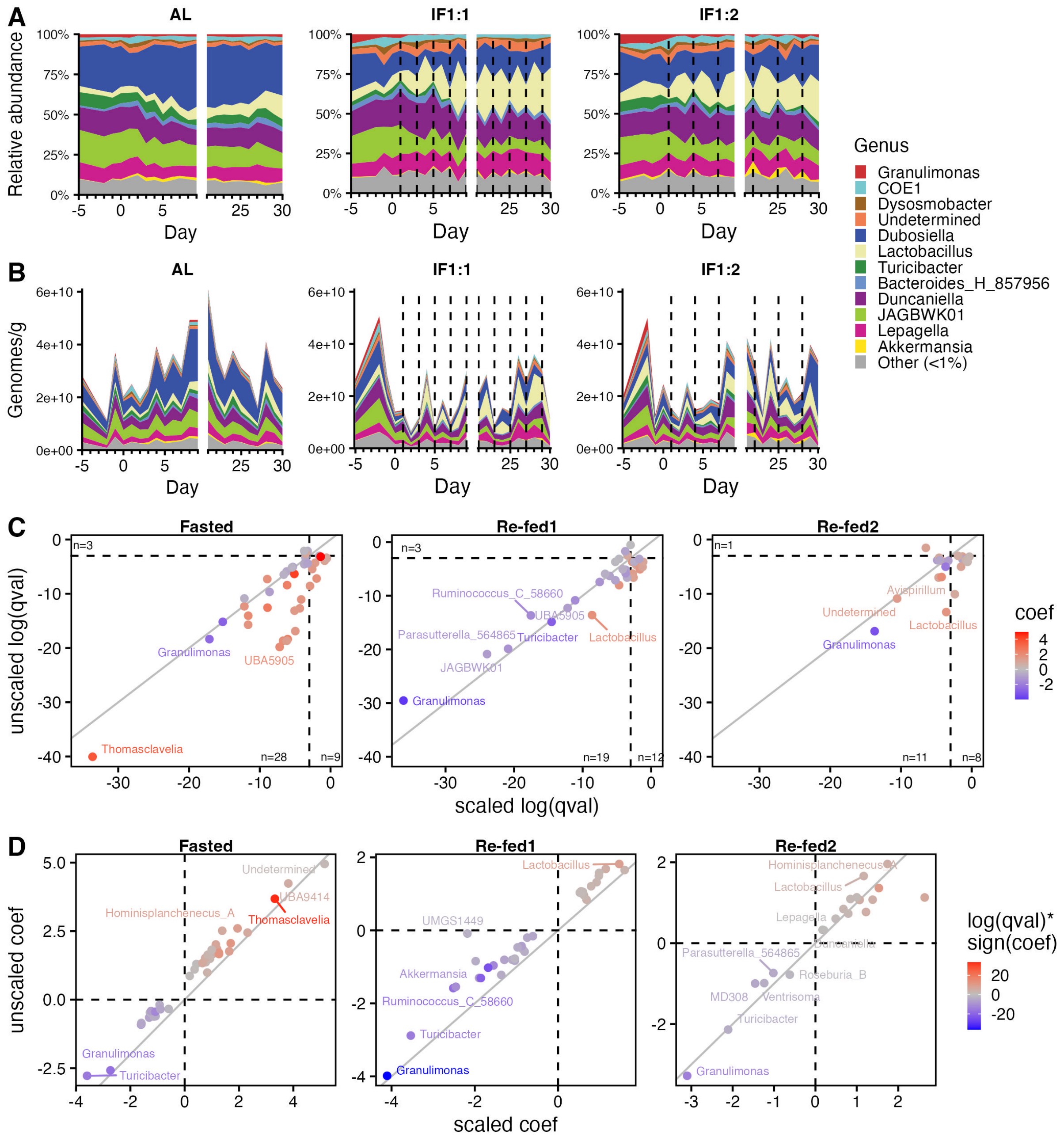


**Figure S2. Impact of absolute bacterial abundance scaling on gut microbiota composition**

(A-B) Mean daily fecal microbiota composition by genus without absolute abundance scaling (A) and scaled to absolute bacterial abundance (B). Fasting days are indicated by dashed black lines.

(C-D) Comparison of differentially abundant taxa significantly associated with feeding status in MaAsLin3 models (fixed effect: feeding status; random effects: mouse ID, source cage ID, day) with or without scaling to absolute bacterial abundance. Diagonal solid grey line marks 1:1 correlation.

(C) q values (FDR-corrected p-values) from scaled versus unscaled models, with significance threshold marked by dashed black lines and counts of genera in each quadrant.

(D) β-coefficients (indicating log2 fold change from baseline/AL fed state) from scaled versus unscaled models, with no effect (coef = 0) marked by dashed black lines.

**Table S2. Output of scaled and unscaled MaAsLin3 models**

Uploaded as separate Excel file

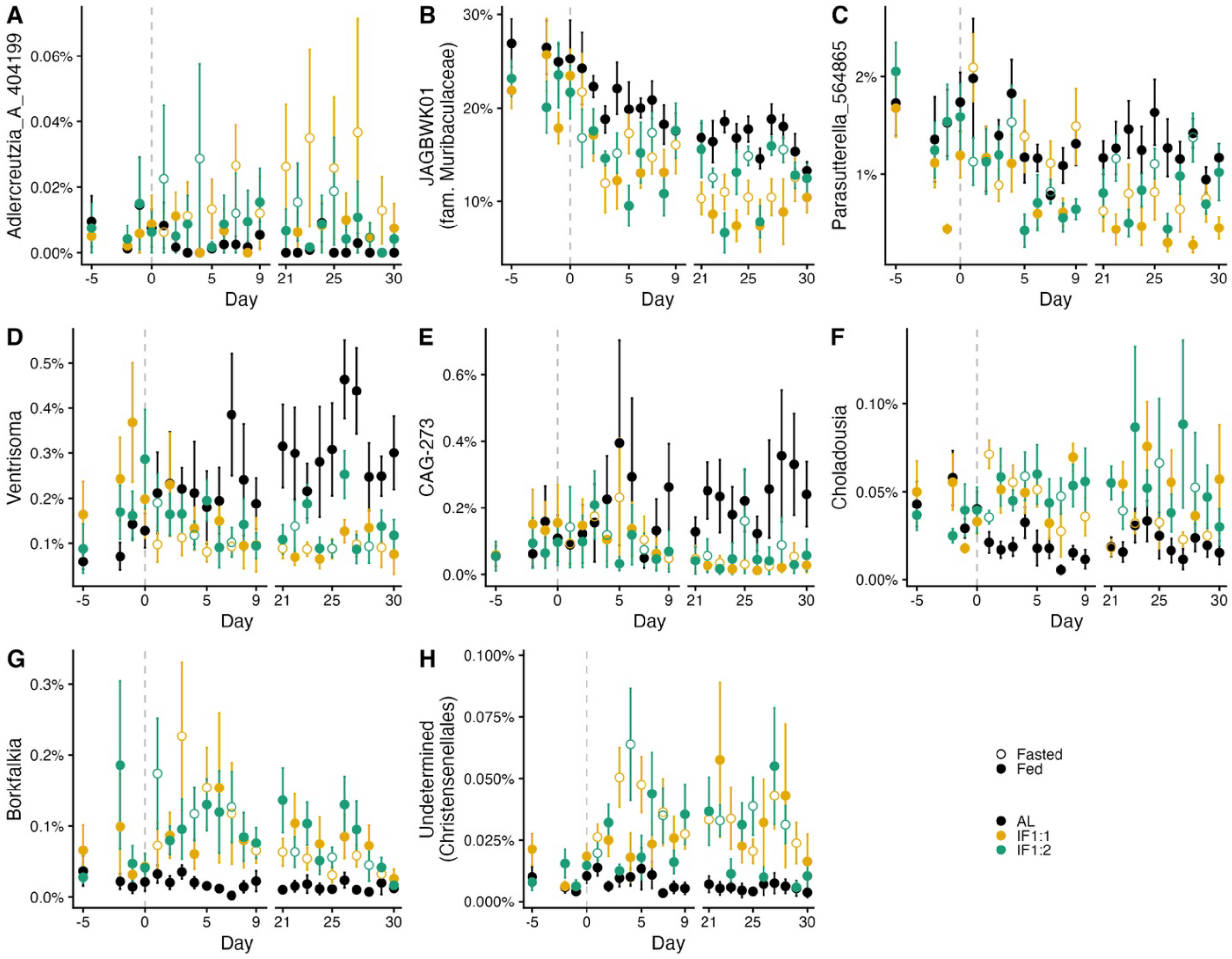


**Figure S3. Gut microbial taxa exhibiting short- and/or long-term responses to intermittent fasting**

Relative abundance of IF-sensitive gut bacterial genera by day of study, with color denoting treatment groups and open and closed circles denoting fasted or fed days, respectively. Mean ± SEM. Identified using MaAsLin3 model (fixed effect: feeding status; random effects: mouse ID, source cage ID, day). Genera exhibited significant changes in relative abundances only during fasting (A) or significant changes in the same direction across both fasted and re-fed days (B-H). See Table S2 for detailed model statistics.
